# Bacteria associated with jellyfish during bloom and post-bloom periods

**DOI:** 10.1101/329524

**Authors:** Maja Kos Kramar, Tinkara Tinta, Davor Lučić, Alenka Malej, Valentina Turk

## Abstract

This study is the first to investigate bacterial community associated with live medusa *Aurelia sp*. in the Gulf of Trieste (northern Adriatic Sea) using both culture independent and culture-based methods. We have analysed bacterial community composition of different body parts of medusa: exumbrella surface, oral arms (‘outer’ body parts) and of gastric cavity (‘inner’ body part) and investigated possible differences in medusa associated bacterial community structure at the time of jellyfish population peak and during senescent phase at the end of bloom, when jellyfish start to decay. Based on 16S rRNA clone libraries and denaturing gradient gel electrophoresis (DGGE) analysis, we demonstrated significant difference between bacterial community associated with *Aurelia* and the ambient seawater bacterial assemblage. Comparing bacterial community composition between different *Aurelia* medusa body parts, communities differed significantly, especially the one within the gastral cavity. The pronounced difference is dominance of *Betaproteobacteria* (*Burkholderia, Cupriavidus* and *Achromobacter*) in gastral cavity of medusa and *Alpha*- (*Phaeobacter, Ruegeria*) and *Gamma*- *proteobacteria* (*Stenotrophomonas, Alteromonas, Pseudoalteromonas* and *Vibrio*) on ‘outer’ body parts. This suggests that body-part specific bacterial association might have an important functional roles for the host. The results of bacterial isolates showed the dominance of *Gammaproeteobacteria*, especially *Vibrio* and *Pseudoalteromonas* in all body parts. Finally, comparison of medusa associated bacterial community structure, at the time of jellyfish population peak and during senescent phase at the end of bloom showed increased abundance of *Gammaproteobacteria*, especially *Vibrio*. Our results suggest members of *Vibrio* group are possible commensal opportunistic visitors, later becoming consumer of moribund jellyfish biomass and that the structure of jellyfish bacterial community might be affected by anthropogenic pollution in the marine environment.

## Introduction

Surfaces of marine animals were found to be a unique habitat for colonization by microorganisms, and the microbial communities associated with living surfaces showed a pronounced variety [1]. Till recently, studies focused on the colonization of benthic organisms such as sponges [2–6], bryozoans [7], and cnidarians, within which are included mainly corals [8–13]. Recent studies of bacteria colonizing crustacean surfaces in the marine pelagic environment, showed considerable dissimilarities with bacterial communities in the surrounding seawater [14–16]. Recently, associated bacteria were reported for gelatinous plankton such as appendicularians [17], ctenophores [18–21], and also cnidarian jellyfish [18,22–28]. Moreover, several studies investigated the role of microbes during jellyfish blooms, and demonstrated high bacterial growth, changes in bacterial community structure in the surroundings of live or decaying jellyfish, and subsequently consequences in altering trophic interactions with higher trophic levels and implications for the carbon, nitrogen, and phosphorus cycles [29–37].

However, very few studies focused on microbial associations with scyphozoan jellyfish during their life span. Studies show a presence of endobiotic bacteria in jellyfish tentacles [28] and suggest that jellyfish could be vectors of bacterial pathogens and implicated in infections of farmed salmons [24,25]. Cleary *et al*. [22] presented data on the bacterial community composition associated with scyphozoan *Mastigias* cf. *papua etpisoni* and box jellyfish *Tripedalia cf. cystophora,* while Weiland-Bräuer *et al*. [27] and Daley *et al*. [18] focused on *Aurelia aurita* s.l. bacterial associates. These studies showed a diverse and specific associated bacterial community, which in composition differs among different marine ecosystems/different jellyfish populations, and has little similarity to the surrounding seawater.

Furthermore, Weiland-Bräuer *et al*. [27] showed that *Aurelia aurita* harbours a different bacterial community on its outer, mucus-covered surface of the exumbrella and gastral cavity, and that microbial community composition differs at different life stages, especially between benthic (polyps and strobila) and sequential planktonic life stages (ephira and juvenile and adult medusa). In addition, an intracellular *Mycoplasma* strain, a possible endosymbiont, has been detected. Studying microbiomes in the gastral cavity of *Cotylorhiza tuberculata, Mycoplasma*-like bacteria was one of four bacterial taxa composing a community of reduced diversity [24]. Most of the bacteria were suggested to have an intracellular lifestyle and established a cooperative relationship with their host. In addition, a new candidate bacterial taxa were proposed [23,26].

Bacterial colonization of given surface is determined by the availability of nutrients, host immune responses, and competition between bacteria from the surrounding environment for attachment space [38]. The epidermis and gastrodermis of jellyfish, including *A. aurita*, contain numerous types of unicellular mucus producing gland cells, leading to the formation of thin, constantly renewing mucus layers over external medusa [39,40]. Under certain conditions like stress, during reproduction and digestion, and also when dying, the amount of released mucus is even more pronounced ([40] and the references within). Mucus on jellyfish surfaces was also found to have a role in surface cleaning and defense against predators. Shanks and Graham [41] characterized mucus secretion as an important chemical defense mechanism, since it contained toxins and discharged and undischarged nematocysts. The contribution to jellyfish chemical defense is, besides mucus, the production of toxins or antimicrobial compounds, such as isolated antibacterial peptide aurelin from mesoglea of *Aurelia aurita* [42].

Further, secreted mucus, whether still covering the surface of jellyfish or already dissociated in forms of blobs, is an attractive niche for bacteria. Since jellyfish mucus is composed mainly of proteins, lipids, and lower percentage of carbohydrates [43], it is a high quality energy source which is readily utilized by bacteria, especially those with a competitive advantage and specialized for settling from surrounding seawater. This indicates that jellyfish as a host can actively or passively affect/select bacterial associates. In addition, bacterial community structure can be also influenced by bacterium-bacterium antagonism, as seen on particles [44], and by environmental conditions determining the presence of metabolically active bacteria and physiological responses of the host [45]. Whether bacteria directly adhere to external cell layers of jellyfish, or are only associated in the thin mucus layer is not clear, however all of the above indicates that the association of bacteria with jellyfish is highly dynamic and complex.

*Aurelia* are among the most widespread scyphozoan medusae that form large aggregations in coastal areas, fjords, and estuaries, and other semi-enclosed or enclosed systems. Moon jelly, *Aurelia* sp. 8 [46] recently designated as *A. solida* [47], is also a very common jellyfish in the northern Adriatic Sea, where 200 years of data show the stabilization of its massive reoccurrence after 2002 [48]. Medusae are generally present from February until late June [49], with peak abundance in the spring [48,50]. The jellyfish outbreaks worldwide seem to become more frequent and last longer in recent years [51]. Whether this is just a rising phase of a natural pattern of decadal oscillations or a true increase of gelatinous zooplankton blooms is still unclear [52]. Still, some data show more frequent and abundant jellyfish aggregations in some coastal areas around the world [48,53], causing numerous socio-economic and ecological problems [54,55]. It has been hypothesized that jellyfish benefit from human-caused changes in environment such as climate change, overfishing, eutrophication, habitat modification, and species introductions [56–59].

This study is the first to investigate the associations of bacteria with live moon jellyfish in the Gulf of Trieste (northern Adriatic Sea) using both culture-independent and culture-based methods. Our hypotheses were the following: (i) the bacterial community associated with medusa is specific and significantly different from the ambient bacterial population in the environment; (ii) the bacterial community composition of different body parts of medusa, i.e. exumbrella surface, oral arms, and of gastral cavity vary; and (iii) medusa-associated bacterial community structure at the time of jellyfish population peak and during senescent phase at the end of bloom, when jellyfish start to decay, differ.

## Materials and Methods

### Sampling site and sampling

The Gulf of Trieste is the northernmost part of the Adriatic Sea. It is characterized by a shallow water column, with salinity and temperature variations, and strong seasonal stratification in late summer [60]. In such an environment, *Aurelia* populations show clear seasonality with late-autumn-early winter recruitment of ephyrae from attached polyps, spring medusa growth, and their decay at high early summer temperatures [59].

Sampling of *Aurelia* medusae was performed during the warmer part of the year in the beginning of May and late June 2011, at the time of adult medusa biomass accumulation. While in May, at the time of population peak, jellyfish were viable and swimming actively, in June, at the end of blooming period, sampled jellyfish were already in the senescent phase and started to decay. Jellyfish were sampled individually by divers or from a boat with a sample bucket. Each individual was placed in plastic bag with some seawater and was transported to the laboratory. Before further analysis, each jellyfish was measured and rinsed twice with sterile seawater (0.2 μm pre-filtered and autoclaved). For determination of the total bacterial community associated with *Aurelia*, samples of exumbrella and oral arms of about 8 cm^2^ in size, were cut out with a sterile razor blade and stored at −80 °C. At the same time, mucus from gastral cavity was sampled with a sterile syringe and stored under the same conditions. At the same time of medusa sampling, ambient seawater samples were collected with a Niskin sampler (V= 5L) at 5 m depth at the oceanographic buoy Vida (45° 32’ 55. 68” N, 13°33’ 1.89” E), where most of jellyfish were restrained at the time of sampling. Each time before sampling standard physical properties, including seawater temperature, salinity and oxygen concentration were measured with a CTD fine-scale probe (Microstructure Profiler MSS90, Sea & Sun Technology GmbH).

### Bacterial isolates from jellyfish and seawater samples

Viable bacterial cells from the surfaces of jellyfish and seawater samples were determined with the spread plate method on modified ZoBell marine agar media [61]. For jellyfish exumbrella, the whole exumbrella surface was inoculated on the plate to create jellyfish imprints of exumbrella-associated bacteria, while the gastro vascular cavity was scraped with a sterile cotton swab and spread evenly over the surface of agar plates. For seawater samples, 100 μL was evenly spread on an agar plate. Inoculated plates were incubated in the dark at an *in situ* temperature for 21 days. For each plate, the number of colony forming units (CFU) was determined and distinctive morphological types of colonies were described. Since direct prints of jellyfish exumbrella were created on agar plates, we estimated the abundance of bacteria as CFU/cm^2^. For DNA extraction, individual colonies were aseptically picked and streaked onto fresh agar plate until single colonies were obtained. A single colony of each bacterial isolate was inoculated in modified liquid ZoBell media and incubated in the dark at the *in situ* temperature until growth was observed (increased turbidity). Part of the liquid cultures was stored in 30% glycerol (final concentration) at – 80 °C for bacterial culture collection. Data on the seawater cultivable bacterial community from the Gulf of Trieste were taken from the dataset gathered during two-year sampling campaign of cultivable bacterial community, of which the sampling time and location coincided with the sampling time of jellyfish (May and June 2011) (Acc. No. KC307273- KC307520). Also, for this dataset sampling was performed at 5 m depth and the bacterial culture collection was obtained as described above.

#### DNA extraction and PCR

Bacterial DNA was extracted from a liquid culture with a modified Chelex-based procedure [62], or with a commercial kit (NucleoSpin Tissue, Macherey - Nagel) according to manufacturer’s protocol. Bacterial cells were harvested by centrifugation and washed twice with 1x PBS buffer (stock 10x; 1.4 M NaCl, 27 mM KCl, 101 mM Na_2_HPO_4_, 18 mM KH_2_PO). When using Chelex (BioRad), the washed cells were re-suspended in 200 μl of 5% Chelex solution. The suspension was incubated for 15 min at 99 °C and transferred at 4 °C for 10 min., and was centrifuged at 4000 rpm for 10 min. The water phase with dissolved DNA was transferred to a new tube and stored at −20 °C until further downstream applications. Bacterial 16S rRNA genes were amplified using universal bacterial primers 27F and 1492R. The PCR reaction mix (50 μl) contained 2 μl (50-100 ng) of extracted genomic DNA, 1x reaction buffer (TrisKCl-MgCl_2_), 2 mM MgCl_2_, 0.2 mM dNTP, 1 μM of each primer, and Taq polymerase (5U/μl, Fermentas). The PCR temperature cycling conditions were as follows: initial denaturation at 94°C for 2 min.; 30 cycles of denaturation at 94°C for 1 min., annealing at 55°C for 2 min., and elongation at 72°C for 2 min. The final cycle was followed by extension at 72°C for 5 min. The quality and size of PCR products was confirmed by agarose gel electrophoresis (1% agarose (Sigma) in 1x TAE buffer) with etidium bromide (10 mg/ml) and visualized using an UV transilluminator (BioDocAnalyze Gel documentation system, Biometra). The bacterial 16S rRNA genes were partially sequenced with 27F primer at Macrogen Inc.

### Total bacterial community composition

#### Jellyfish-associated bacterial community DNA extraction

Exumbrella and oral arms samples were thawed down. Bacterial DNA was extracted with CTAB (cetyl-trimethyl-ammonium bromide) as described before [21] with slight modification. Samples were placed into a tube containing 2% CTAB solution (1.4 M NaCl, 100 mM Tris_Cl pH= 8, 2% CTAB, 20 mM EDTA pH=8, 0.2% β-mercaptoethanol), and were incubated at 65°C for 1 h. Afterwards, a STE buffer (6.7% sucrose, 50 mM Tris-Cl pH= 8, 1 mM EDTA pH= 8) and proteinase K (100 μg/ml final concentration, Sigma) were added and the mixture was incubated at 55°C overnight. Following chloroform-isoamylacohol (24:1, v/v, Sigma) and phenol- hloroform- isoamylacohol (25:24:1, v/v, Sigma) purification steps, DNA was precipitated at - 20°C overnight with isopropanol. The pellet was washed with 70% ice-cold ethanol and dried in a speed vac. The precipitated DNA was re-suspended in 0.22 μm pre-filtered, autoclaved 1X TE buffer, and kept at – 20°C.

#### Seawater’s total bacterial community DNA extraction

Seawater samples were filtered onto 0.2 μm polyethersulfone membrane filters (47 mm diameter, PALL Inc.), which were stored at - 80°C. DNA was extracted from the filters (one quarter per sample) as described in Böstrom *et al*. [63], with slight modifications. DNA was precipitated at - 20°C for 1 h, with 0.1 volume of sodium acetate (3 M NaAc, pH= 5.2) and 0.6 volume of isopropanol. The pellet was washed with 70% ice-cold ethanol and dried in a speed vac. Precipitated DNA was re-suspended in 0.22 μm, pre-filtered, autoclaved TE buffer, and kept at - 20°C.

#### Denaturing Gradient Gel Electrophoresis

For DGGE analysis of jellyfish-associated and seawater’s total bacterial community, the bacterial 16S rRNA genes were amplified using a universal primer set, 341F with a 40 bp GC-clamp and 907R as described before [64,65]. The PCR reaction mix with a final volume 50 μl contained 2 μl of extracted DNA (50–100 ng), 1x reaction buffer (Tris KCl-MgCl_2,_ Fermentas), 1.5 mM MgCl_2_ (Fermentas), 0.2 mM dNTP (Fermentas), 0.5 μM of each primer (Sigma), 0.38 μg/ml BSA (Fermentas), and Taq polymerase (5 U/μl, Fermentas). The PCR touchdown protocol according to Don *et al*. [66] was used: with initial denaturation at 94°C for 5 min., followed by 10 touchdown cycles and 20 standard cycles: denaturation for 1 min. at 94°C, primer annealing for 1 min. at 55°C, and primer extension for 3 min. at 72°C. The last cycle was followed by 2 min. incubation at primer extension temperature of 72°C.

When we were unable to obtain a sufficient quantity of PCR products from jellyfish samples, we used a two-step nested PCR- DGGE strategy [67], modified to analyze the marine bacterial community. Bacterial 16S rRNA genes were first amplified with universal primer set, 27F and 1492R. The PCR reaction mix, with a final volume 50 μl, contained 2 μl of extracted DNA (50-100 ng), and was prepared the same as described above in this section. The PCR temperature cycling conditions were as follows: initial denaturation for 2 min. at 94°C, followed by 25 standard cycles: denaturation at 94°C for 1 min., primer annealing for 1 min. at 50°C, and primer extension at 72°C for 1 min. The last cycle was followed by 5 min. incubation at the primer extension temperature of 72°C. Second, nested amplification was performed using a DGGE primer set, PCR mixture, and a touchdown annealing protocol, as described above in this section. The quality and size of PCR products were tested by agarose gel electrophoresis. PCR products were analyzed by DGGE electrophoresis, as previously described in [33].

Distinct bands were excised from the gel and placed in 100 μl of sterile Sigma water overnight to elute DNA. The eluted DNA was re-amplified using primer set 341F and 907R and the same reaction mix (a final volume 50 μl) with 2 μl of eluted DNA, as described above in the first paragraph in this section. The cycling protocol used was the same as to amplify the DNA of bacterial isolates (see section on Bacterial isolates DNA extraction and PCR). The bacterial 16S rRNA genes were partially sequenced with 341F primer at Macrogen Inc.

#### Bacterial 16S rRNA gene clone libraries

For jellyfish and seawater samples clone libraries construction, bacterial 16S rRNA gens were amplified using the same DNA as for DGGE and universal primer set, 27F and 1492R, as described before [33]. The PCR reaction mix with a final volume 50 μl contained 2 μl of extracted DNA (50-100 ng), and was prepared as described above (see section on DGGE, first paragraph). For samples with low DNA concentration (extracted from jellyfish samples), a nested PCR-libraries approach was used [68], and modified to analyze marine bacterial community. Again, bacterial 16S rRNA gene was first amplified with a universal primer set, 27F and 1492R using same protocol and reaction mix as in the first amplification step of the nested PCR-DGGE strategy (see the section on DGGE, second paragraph). Second, nested amplification was performed using primers 341F and 907R. The PCR reaction mixture and cycling protocol were the same as used for clone library construction (described above in this section [33]). The PCR products were immediately ligated into a commercially available pCR 2.1 vector and transformed into competent *E. coli* TOP 10 cells using a commercially available TA Cloning Kit (Invitrogen), and according to the manufacturer’s protocol. The plasmid inserts from of each clone library were partially sequenced using M13F primer or 27F primer at Macrogen Inc.

### Sequence analyses

Raw sequence data recovered from bacterial isolates and clone libraries were passed through the DNA Baser program (www.DNAbaser.com) to remove traces of sequencing primer, and to trim away ambiguous bases at the end of a sequence. The clone libraries sequences were also checked for vector contamination and were analyzed with the program Bellerophon (https://greengenes.lbl.gov/) to detect chimeric sequences, which were removed. Additionally, Mothur software [69] was used to further reduce poor quality sequence data. Sequence taxonomic identities (with ≥ 97%) of bacterial isolates and sequences recovered from clone libraries and DGGE bands were assigned using the BLAST (Basic Local Alignment Tool) algorithm available at NCBI (National Centre for Biotechnology Information). Around 50% of the sequences recovered from clone libraries and DGGE bands exhibited <97% similarity to previously published GenBank entries (omitted from further analysis). Classification of bacterial isolates was done down to the genus level, and of clones and DGGE bands down to the family level. The number (N) of high quality sequences, with ≥ 97% similarity to the nearest GenBank entry, and the number of distinct bacterial genus and families (S_obs_) is presented in Supporting Information (S1 Table, S2 Table). The contribution of distinct bacterial genus or families was expressed as a percentage of the total number of sequences in each sample or library (relative abundance) (S1 Table, S2 Table). Chloroplast sequences were omitted from further analysis.

#### Nucleotide sequence accession numbers

The 16S rRNA gene sequences, for all bacterial isolates, clone libraries, and DGGE bands, obtained in this study have been deposited in the GenBank (NCBI) under following accession numbers: from KF816449 to KF816471, and KF816480 to KF816592 for bacterial isolates (Supporting information, Table S5), from KF816761 to KF816832, from KF817469 to KF817519, from MF952738 to MF952748, and from MF952764 to MF952865 for sequences obtained from clone libraries, and from MF952749 to MF952763 for sequences obtained from DGGE bands.

### Diversity indices and statistical analyses

To compare the diversity of viable bacterial isolates from jellyfish and surrounding seawater, ecological diversity indices were calculated for each sample: the number of different bacterial genus (species richness (Sobs)), Shannon diversity index (H’), Margalef’s index (d), Pielou’s evenness index (J’) and Chao-1 index. The same parameters were calculated for 16S rRNA bacterial clone libraries at the family level. Additionally, in order to estimate how well the actual species composition was captured, for each clone library a coverage value was calculated as C= 1-n_1_/N, where n1 is the number of phylotypes appearing only once in the library, and N is the library size [70].

Non-metric multi-dimensional scaling (nMDS) plots were used to determine the similarities between DGGE banding patterns. For this purpose, a similarity matrix was calculated (using Jaccard resemblance measure) based on the presence/absence matrix of align bands. Analysis of similarity (ANOSIM) was used to verify the significance of similarity among bacterial communities, as indicated by nMDS, by testing the hypothesis that bacterial communities from the same cluster are more similar to each other than to communities in different clusters. Cluster analysis was used to determine scaled similarities between 16S rRNA gene clone libraries (total bacterial communities) and between bacterial isolates (culturable bacterial communities). For cluster analysis of 16S rRNA gene clone libraries, a Bray-Curtis similarity matrix was constructed from arcsine-transformed relative abundances of distinct bacterial families in each clone library. For bacterial isolates, a Bray-Curtis similarity matrix was constructed from untransformed relative abundances of distinct bacterial genus in each culturable bacterial community. Based on the similarity matrix, dendrogram was produced with group-average linkage algorithm. The similarity profile test (SIMPROF) was used to define statistically significant clusters in samples.

To examine the difference between communities associated to different jellyfish body parts and seawater, one-way ANOSIM statistic with 999 permutations, based on Bray-Curtis similarity matrix, was made. Samples were grouped according to isolation source (communities of jellyfish exumbrella (AK), jellyfish oral arms (AR) and jellyfish gastral cavity (AG) and communities of seawater (W)). Similarly, one-way ANOSIM statistic with 999 permutations was made to examine the difference between communities associated with jellyfish at the time of population peak and at the end of blooming period. Samples were grouped according to time (communities associated to jellyfish sampled at time of population peak (May) and those sampled at the end of the bloom, when jellyfish were in senescent phase (June)). Additionally, similarities percentage (SIMPER) analysis was used to determine which bacterial group contribute the most to the differences between communities of different body parts of jellyfish and water communities (for culturable and total bacterial community). Diversity indices and statistical analysis were performed using Primer v6 [71] and PAST, version 3.9 [72].

## Results

The composition of the bacterial community associated with scyphomedusae *Aurelia,* which frequently blooms in the Northern Adriatic, was studied and compared with the community composition from the surrounding seawater, in order to understand if the jellyfish-associated community is specific and significantly distinct from the ambient seawater bacterial assemblage. The composition of bacterial community associated with different jellyfish compartments (exumbrella surface, oral arms, and in the gastral cavity) was analysed to examine the compartment-specificity of associated bacterial consortia. In addition, we compared the composition of the bacterial community associated with jellyfish collected during two different time points of bloom development/progression: (i) at the peak of population, and (ii) at the end of the blooming period/at the decay of the bloom. The bacterial community composition/structure was determined using both culture-independent and culture-dependent techniques.

### Comparison of jellyfish-associated and ambient seawater bacterial community composition

The phylogenetic analyses of the bacterial 16S rRNA gene clone libraries revealed significant difference between jellyfish-associated and ambient seawater bacterial communities (ANOSIM, global R= 0.777, p< 0.01) (Fig 1A). The bacterial communities associated with *Aurelia* showed the dominance of bacterial phyla *Proteobacteria*, which consisted of *Alphaproteobacteria* (up to 75%), *Gammaproteobacteria* (up to 45.5%), and *Betaproteobacteria* (up to 53.5%), with different relative contributions in the individual jellyfish sample (Fig 1B). At the family level, *Alphaproteobacteria* were dominated by *Rhodobacteraceae* (mostly *Phaeobacter, Ruegeria*)) and *Betaproteobacteria by Burkholderiaceae* (*Burkholderia*) (Fig 1B, S1 Table). Within *Gammaproteobacteria,* mostly *Vibrionaceae* (*Vibrio*), *Pseudoalteromonadaceae* (*Pseudoalteromonas*), *Xanthomonadaceae* (*Stenotrophomonas*), and *Pseudomonadaceae* (*Pseudomonas*) (Fig 1B, S1 Table) were detected.

**Fig 1. Bacterial 16S rRNA gene clone libraries constructed from samples of *Aurelia* jellyfish and ambient sweater.**

(**A**) Cluster analysis based on bacterial 16S rRNA gene clone libraries. AK-jellyfish exumbrella surface, AR-jellyfish oral arms, AG-mucus from gastral cavity and W-ambient seawater. Samples were collected in May (grey squares) and June (inverted black triangles).The dendrogram was inferred with the group average algorithm, based on the Bray– Curtis similarity matrix of arcsine transformed averaged abundances. Grey branches do not differ significantly (SIMPROF test, p> 0.05). (**B**) Composition of bacterial 16S rRNA gene clone libraries (expressed as percentage of clones) constructed from samples of jellyfish exumbrella surface (AK1, AK2), oral arms (AR1), mucus from gastral cavity (AG1) and the ambient seawater (W_May) sampled in May and jellyfish exumbrella surface (AK6, AK7), oral arms (AR6) and the ambient seawater (W_Jun) sampled in June. Cumulative column charts represent relative abundances of bacterial family and area chart in the background represent relative abundances of major bacterial groups and *Proteobacteria* class. Taxa with relative abundance of< 3% across all samples are subsumed under Other Bacteria.

In comparison to the jellyfish-associated bacterial community, the ambient seawater bacterial community was more diverse and dominated by three bacterial phyla: *Proteobacteria, Flavobacteria*, and Cyanobacteria (Fig 1B, S5 Table). *Alphaproteobacteria* (up to 38.6%) were dominated by *Rhodobacteraceae* and SAR11, *Gammaproteobacteria* (up to 21.4%) by *Litoricolaceae* and SAR86; *Flavobacteria* (up to 17.4%) by *Flavobacteriaceae* and *Cryomorphaceae*, and *Cyanobacteria* (up to 15.4%) by *Synechococcus*. We also detected *Actinobacteria* (10.3% in May) with the representative from the *Microbacteriaceae* family. (Fig 1B, S1 Table). According to SIMPER analysis *Synechococcus*, SAR11 and *Flavobacteriaceae* contributed the most to difference between jellyfish-associated and water column bacterial community (S3 Table).

### The bacterial community composition associated with different body parts of jellyfish

The results of 16S rRNA gene clone libraries analysis pointed on the statistically significant differences between bacterial communities associated with different body parts of jellyfish (exumbrella surface, oral arms, and gastral cavity) (ANOSIM, global R= 0.571, p< 0.05)(Fig 1A). The bacterial communities’ composition associated with different body parts of jellyfish sampled at the peak of population, were as follows. The bacterial communities associated with jellyfish exumbrella were dominated by *Alphaproteobacteria* (up to 75%), followed by *Gammaproteobacteria* (up to 22.2%) and *Betaproteobacteria* (up to 12.5%) (Fig 1B). The population of *Alphaproteobacteria* was dominated by *Rhodobacteraceae*, mostly *Phaeobacter, Ruegeria*, but also *Rhizobiaceae, Hyphomicrobiaceae*, and *Sphingomonadaceae* were detected. Within *Gammaproteobacteria*, mostly *Xanthomonadaceae* (*Stenotrophomonas*), but also *Alteromonadaceae* (*Alteromonas*), and within *Betaproteobacteria* exclusively *Comamonadaceae* were detected. (Fig 1B, S1 Table). The bacterial community of jellyfish oral arms was more diverse than the bacterial community associated with exumbrella and gastral cavity (S5 Table). The bacterial community associated with oral arms consisted of *Alphaproteobacteria* (50%, exclusively *Rhodobacteriaceae*) and a higher percentage of *Gammaproteobacteria* (31.3%) composed mainly of *Vibrionaceae* (*Vibrio*), but also *Pseudoalteromonadaceae, Moraxellaceae*, and *Pseudomonadaceae. Betaproteobacteria* were detected (12.5%, only *Burkholderiaceae*), and also a small percentage of *Actinobacteria* (6.3%) (Fig 1B, S1 Table). In contrast, the bacterial community in the gastral cavity, was dominated by *Betaproteobacteria* (53.5%), followed by *Gammaproteobacteria* (27.9%) and *Actinobacteria* (11.6%, dominated by *Micrococcaceae*). At the family level, *Burkholderiaceae* (*Burkholderia*) and *Alcaligenaceae* dominated the *Betaproteobacteria* class. The gammaproteobacterial population was almost exclusively *Pseudomonadaceae* (*Pseudomonas*) (Fig 1B, S1 Table).

The bacterial community structure of different jellyfish body parts was also studied using denaturing gradient gel electrophoresis (DGGE). Bacterial community fingerprints varied, both within and between sample types (exumbrella, oral arms, and gastral cavity). Despite the observed heterogeneity (S1 Fig), the DGGE-based non-metric multidimensional scaling (nMDS) analysis showed that bacterial communities clustered according to jellyfish body part (S1 Fig) (ANOSIM, global R= 0.633, p< 0.05). Jellyfish-associated bacterial community composition was more similar between replicates of the samples collected from the same body parts (40%) than different jellyfish body parts (Fig 2). Phylogenetic information obtained from excised DGGE bands showed that bacterial taxa across all samples mostly belonged to *Alphaproteobacteria* (*Roseobacter, Phaeobacter, Ruegeria* all *Rhodobacteraceae*), but also *Gammaproteobacteria* (*Vibrio, Pseudoalteromonas, Stenotrophomonas*) and *Betaproteobacteria* (*Burkholderia*) (S1 Fig, S7 Table).

Fig 2. **Non-metric multidimensional (nMDS) analysis based on bacterial community DGGE banding patterns of *Aurelia* jellyfish samples.**

AK- jellyfish exumbrella surface, AR-jellyfish oral arms, AG- mucus from gastral cavity and W- ambient seawater. Samples were collected in May (grey squares) and June (inverted black triangles). Resemblance circles: grey line - 40% similarity; black line - 50% similarity.

### Bacterial community structure shifts due to jellyfish population senescence

Our results show the difference between the bacterial communities associated with jellyfish collected during the peak of the jellyfish bloom and one month later, at the jellyfish population senescence (Fig 1). Changes in bacterial communities, due to jellyfish population senescence were evident as the shift towards *Gammaproteobacteria,* mostly at the expense of *Betaproteobacteria*, and to a lesser extent at the expense of *Alphaproteobacteria*, whose dominance became less pronounced. In addition, diversity was lower in the bacterial community associated with senescent jellyfish S5 Table). Bacterial communities associated with exumbrella surface were composed of *Gammaproteobacteria* (up to 42.9%) and *Alphaproteobacteria* (up to 66.7%; almost exclusively *Rhodobacteriaceae* of which mostly *Roseovarius, Ruegeria*). Within *Gammaproteobacteria*, previously dominant *Xanthomonadaceae* were replaced with *Alteromonadaceae* (*Marinobacter*) and *Vibrionaceae* (*Vibrio*). In addition, *Pseudoalteromonadaceae* (*Pseudoalteromonas*) and *Moraxellaceae* were detected (Fig 1B, S1 Table). A similar change/shift was evident in the bacterial community structure determined in the sample of oral arms. *Gammaproteobacteria* were dominant (45.5%; taxa composition similar to that associated with exumbrella surface, with even more pronounced *Vibrio* dominance), followed by *Alphaproteobacteria* (36.4%, exclusively *Rhodobacteraceae*) and *Betaproteobacteria* (18.2%, exclusively *Burkholderiaceae*). The shift in bacterial community structure was supported by SIMPER analysis showing that *Rhodobacteriaceae* and *Comamonadaceae* were more frequent in the bacterial community associated with jellyfish at the peak of the bloom, and *Rhodobacteriaceae, Vibrionaceae*, and *Alteromonadaceae* in the bacterial community associated with senescent jellyfish (S8 Table).

### Culturable bacterial community composition

Altogether, 135 bacteria were isolated from the exumbrella surface and gastral cavity of jellyfish, sampled during the peak of jellyfish population (AK1, AK3, AK6, AG1, AG6), and at the end of the blooming period (AK8, AK10, AK11, AG8, AG11). Regarding the morphology of bacterial colonies, we detected nine morphotypes in May, and only three morphotypes in June. The estimated abundance of cultural bacteria associated to jellyfish exumbrella was 1.9 CFU/cm^2^ in May, and 2.0 CFU/cm^2^ in June. Identification of 16S rRNA gene sequence revealed that across all samples, bacteria predominantly belonged to *Proteobacteria* (*Gammaproteobacteria* followed by *Alphaproteobacteria and Betaproteobacteria*), *Actinobacteria,* and *Firmicutes* (*Bacilli)* (exhibiting ≥ 97% identity to previously described bacterial species). Analysis of bacterial isolates again showed significant differences between seawater and jellyfish-associated communities (ANOSIM, global R=0.393, p< 0.05) (Fig 3), with seawater communities being more diverse (Fig 4, S6 Table). *Gammaproteobacteria* (mostly *Vibrio* and *Pseudoalteromonas*) dominated the exumbrella surface-associated community (up to 100%) and the community of gastral cavity (> 60%), while bacterial isolates obtained from ambient seawater were mainly affiliated with *Alphaproteobcateria* (88% in May and 42% in June, mostly *Erythrobacter* (*Erythrobacteraceae*) and *Brevundimonas* (*Caulobacteraceae*)), followed by *Gammaproteobacteria* (32%, mostly *Halomonas* (*Halomonadaceae*), *Idiomarina* (*Idiomarinaceae*)) and *Actinobacteria* (25%, *Brevibacterium* (*Brevibacteriaceae*)) which were more abundant in June (Fig 4, S2 Table). According to SIMPER analysis, *Erythrobacter, Brevibacterium*, and *Brevundimonas* contributed the most to the difference between jellyfish-associated and water communities (S4 Table).

**Fig 3. Cluster analysis based on culturable fraction of bacterial community associated with *Aurelia* jellyfish.**

AK- jellyfish exumbrella surface, AG- mucus from gastral cavity and W- ambient seawater. Samples were collected in May (grey squares) and June (inverted black triangles). The dendrogram was inferred with the group average algorithm, based on the Bray–Curtis similarity matrix of arcsine transformed averaged abundances. Grey branches do not differ significantly (SIMPROF test, p> 0.05).

**Fig 4. Bacterial isolates obtained from *Aurelia* jellyfish and ambient seawater.**

(**A**) Bacterial isolates obtained from jellyfish exumbrella surface (AK1, AK3, AK6), gastral cavity (AG1, AG6) and seawater (W_May) in May in the Gulf of Trieste. (**B**) Bacterial isolates obtained from exumbrella surface (AK8, AK10, AK11) and gastral cavity (AG8, AG11) of jellyfish and seawater (W_Jun) in June in the Gulf of Trieste. Cumulative column charts represent relative abundances of bacterial genus and area chart in the background represent relative abundances of major bacterial groups and *Proteobacteria* class. Taxa with relative abundance of< 3% across all samples are subsumed under Other Bacteria.

Differences between culturable bacterial communities of different body parts were small. Bacterial communities associated with jellyfish exumbrella at the time of population peak were dominated by *Gammaproteobacteria* (from 44.4% up to 100%), while *Alphaproteobacteria* and *Actinobacteria* represented up to 22.2% of the community (Fig 4A). Small percentages of isolates belonged to *Betaproteobacteria* (11%, exclusively *Delftia Comamonadaceae*) *Bacilli* (9%; mostly *Exiguobacterium*) and *Bacteriodetes* (12.5%). Considering the main representatives within *Gammaproteobacteria, Vibrio* (*Vibrionaceae*), *Pseudoalteromonas* (*Pseudoalteromonadaceae*), and *Stenotrophomonas* (Xanthomonadaceae) dominated, but also *Pseudomonas* (*Pseudomonadaceae*), *Alteromonas* (*Alteromonadaceae*), and *Psyhrobacter* (*Moraxellaceae*) were detected. Representatives of *Alphaproteobacteria* were mostly *Labrenzia* and *Phaeobacter* (*Rhodobacteraceace*), and representatives of *Actinobacteia* mostly *Kocuria* (*Micrococcaceae*) and *Microbacterium* (*Microbacteriaceae*) (Fig 4A, S2 Table). The composition of bacterial communities in jellyfish gastral cavity was similarly dominated by *Gammaproteobacteria* (> 60%), followed by *Alphaproteobacteria* (up to 40%) and *Betaproteobacteria* and *Bacilli* (both 10%). The population of *Gammaproteobacteria* was dominated by *Pseudoalteromonas* followed by *Pseudomonas, Vibrio*, and *Stenotrophomonas* (Fig 4A, S2 Table).

Changes in culturable bacterial community composition, presumably related to jellyfish population decay, at the end of blooming period were significant (ANOSIM, global R= 0.362, p< 0.05). Changes were evident as there was even more pronounced dominance of *Gammaproteobacteria* (from 66.7% up to 100%) in communities associated to jellyfish exumbralla and of gastral cavity (Fig 4B). Within the Gammaproteobacterial population, predominant community members *Pseudoalteromonas, Stenotrophomonas*, and *Pseudomonas* almost or completely ‘disappeared’, and *Vibrio* became highly dominant or even formed a monoculture representative (Fig 4B, S2 Table). The change in community structure towards *Vibrio* was also confirmed by SIMPER analysis (S9 Table). Consequently, the prevalence of *Vibrio* also resulted in lower diversity (S6 Table).

## Discussion

### *Aurelia* associated bacterial community

#### Comparison of jellyfish-associated and ambient seawater bacterial community composition

Our results on bacterial community composition, assessed by culture-dependent and cultural-independent approaches, demonstrated significant differences between bacterial community associated with *Aurelia* and the ambient seawater bacterial assemblage. Phylogenetic analysis showed a wide diversity of bacterial community associated with jellyfish, including members of Proteobacteria (*Alpha-, Gamma-*, and *Beta-proteobacteria*), which dominated the community, and members of *Actinobacteria* and *Cyanobacteria*. Ambient seawater bacterial communities were more diverse, and dominated by three bacterial phyla: *Proteobacteria, Flavobacteria*, and *Cyanobacteria*. Also, within Proteobacterial groups associated with jellyfish and the one detected within ambient seawater assemblage, had different taxonomic affiliations, and the dominance or exclusivity of only one taxon was hardly present.

Similar observations of the jellyfish-specific bacterial community, distinct from the community in ambient seawater, were reported previously for *A. aurita* [18,27] and also other marine animals [45]. Since associated bacterial assemblages differed from the ambient seawater bacterial community, and from bacteria associated with other types of substrates/surfaces found in the water column, it was suggested that associations with animals might be specific to some degree [45]. According to Taylor *et al*. [73] sponge bacterial associates could be separated/split into three groups: (i) bacterial specialists - found on only one host species; (ii) host associates - found on multiple hosts; and (iii) generalists - found on multiple hosts and within the seawater community. In our study most bacteria associated with *Aurelia* were not detected in the ambient seawater, however, they were closely related to bacteria previously found in association with other host animals (exhibiting at least 97% similarity), indicating that this relationship is not host-specific. Previous studies on *A. aurita* bacterial associates also did not reveal the presence of any *Aurelia*-bacterial specialists, with the exception of *Mycoplasma* sp. (class *Mollicutes*), a possible/hypothetical endosymbiont [18,27]. However, in our study, we were not able to detect any *Mycoplasma* members. In addition, bacterial community composition was very different to the community associated with *A. aurita* from the North West Atlantic and the Baltic Sea [18,27]. This might suggest the possible effect of host genetics background (different populations of *Aurelia* species in geographically distant locations), and also the important’s of environmental and anthropogenic conditions, determining the presence, activity, and composition of bacterial community in jellyfish’s environment.

We found another interesting result, which is the consistently unsuccessful amplification of bacterial 16S rRNA genes from jellyfish samples, unless an additional nested PCR reaction was performed. Problems with DNA amplification were reported before in the analyses of the tissue of healthy corals, and were attributed to the low abundance of bacterial associates [74], confirming previous observation of rare isolated bacterial cells within coral tissue by in situ hybridization [75]. This indicates that unsuccessful DNA amplification in our study also could be the consequence of the overall low bacterial number on jellyfish exumbrella, the oral arm surface, and in the gastral cavity. The speculations on the low number of *Aurelia* jellyfish-associated bacteria, was also confirmed by scanning electron microscopy of the adult medusa umbrella surface in our parallel study (data not shown). Our results show that the surface was covered with mucus, and we observed mucus secretions in the form of flocs on all external surfaces, but with no bacteria observed. Unlike the epidermal umbrella surface, the examination of mucus secreted from exumbrella surface, revealed the presence of considerable amounts of bacteria (data not shown). Using the same microscopic method, Johnston and Rohwer [76] similarly found that external cell layers of coral are invariably clean of adhering microbes. They did, however, suggest the possibility of a dynamic community hovering in the boundary layers above the coral epidermis. So, the possibility that the majority of bacteria dwell in mucus produced by medusa rather than being present within the mesoglea or attached to epidermis is also in agreement with observations by Weiland-Bräuer *et al*. [27], detecting the majority of bacteria located on the outer surface of coating mucus, covering *A. aurita* polips. This could also support our results of bacterial colonies grown after imprints of jellyfish surfaces on agar plates, where it was estimated CFU less than two bacterial colonies/cm^2^ of jellyfish surface in May and June. The presence of rare bacterial cells could be due to the fact that adult medusa has evolved mechanisms of defense against epibiotic organisms. One type of mechanism could be the production of antibacterial peptide aurelin, extracted from mesoglea of *A. aurita* [42]. Based on that, we can speculate that *A. aurita* is a ‘hostile’ environment for bacteria. It is also known that jellyfish surfaces, including *A. aurita,* are covered by a constantly renewing mucus layer, which was found to have implications in surface cleaning and defense against predators [39–41]. Similarly as Garren and Azam [77] demonstrated for corals, surface cleaning by mucus production could be used by jellyfish to regulate an abundance of bacterial associates. Even more so, during certain conditions, including stress, the mucus production is more pronounced [41]. In our study, extensive mucus release (from surface and gastral cavity) was detected during the processing of jellyfish, possibly leading to bacterial loss and low bacterial numbers of jellyfish associates in our samples.

Despite being rare, the total and the culturable part of the bacterial community associated with *Aurelia* jellyfish from the Gulf of Trieste was found to be diverse. It was composed mostly of: (i) bacteria belonging to genus *Ruegeria, Phaeobacter*, and *Pseudoalteromonas*, known for their extraordinary ability to successfully compete and colonize surfaces, and also to enhance survival chances of the host organism [78–81]; (ii) bacteria belonging to genus *Alteromonas* and *Vibrio*, known as particle and surface colonizers with the ability to degrade and utilize a broad spectrum of organic substrates [81]; and (iii) bacteria belonging to genus *Stenotrophomonas, Burkholderia, and Achromobacter*, mostly known as medically important strains, but also with high bioremediation potential due to ability of PAHs and xenobiotic degradation and with broad antibiotic resistance [82–87]. With the exception of *Ruegeria, Burkholderia, and Achromobacter*, other bacteria were also recovered by culturing. The presence of *Ruegeria* and *Phaeobacter* (*Alphaproteobacteria* - *Roseobacter* clade) and *Pseudoalteromonas, Alteromonas*, and *Vibrio* (*Gammaproteobacteria*) is not surprising, since they are known as successful and dominant particle/surface colonizers [80,88,89], and regularly associated with marine sponges [4], corals (tissue and mucus) [8,9,13,90–93], and ctenophores [18,20,94]. A more interesting feature of *Aurelia* jellyfish associated bacterial community was the high relative abundance of *Burkholderia* and *Achromobacter* (*Betaproteobacteria*). This group is not characteristic of a marine environment but was found in environments characterized with lower salinity and higher nutrient concentrations, such as estuaries [95]. *Betaproteobacteria* were also found in association with sponges [3,5,96], corals [9,11,12], ctenophores [19,94], and cnidarian Hydra [97], but, with the exception of the last, their relative abundances were lower than in our study. Of the above bacterial taxa, only *Phaeobacter, Vibrio,* and *Pseudoalteromonas* were previously found in association with *A. aurita* [18,27,33].

#### The bacterial community composition associated with different body parts of jellyfish

Bacterial community composition differed significantly between different *Aurelia* medusa body parts, especially the one within the gastral cavity. The communities of exumbrella and oral arms shared dominant bacterial groups, *Alphaproteobacteria* followed by *Gammaproteobacteria*, while the community in the gastral cavity was dominated by *Betaproteobacteria*, followed by *Gammaproteobactera* and *Actinobacteria*. Within *Alphaproteobacteria*, bacterial communities of the exumbrella surface and oral arms were affiliated with *Phaeobacter, Ruegeria*, and within *Gammaproteobacteria* with *Stenotrophomonas, Alteromonas, Pseudoalteromonas*, and *Vibrio*. In the gastral cavity were members of *Betaproteobacteria* affiliated with *Burkholderia, Cupriavidus*, and *Achromobacter*. Members of *Gammaproteobacteria* affiliated mostly with *Pseudomonas*, and members of *Actinobacteria* with *Kocuria*.

In contrast to the total bacterial community, bacteria isolated from jellyfish (from both the exumbrella surface and gastral cavity) were mostly affiliated with *Gammaproteobacteria,* within the most relevant members affiliated with *Vibrio, Stenotrophomonas*, and *Pseudoalteromonas*. The observed dominance of different bacterial classes within the total bacterial community and cultured bacterial community in our study is not that surprising, since both culturing and molecular-based methods are biased towards certain microbial groups. Similar observations, were reported previously by Rohwer [11] studying coral-associated bacterial communities. In addition, the culturing approach revealed the presence of bacteria affiliated with *Microbacterium* (*Microbacteriaceae*), *Sphingobacterium* (*Sphingobacteriaceae*), *Brevundimonas* (*Caulobacteraceae*), and *Delftia* (*Comamonadaceae*). However, considering the main representatives within bacterial groups, molecular-based studies and the culture-dependent method more or less pointed to the presence/dominance of the same bacterial taxa.

The pronounced difference in composition between gastral cavity bacterial community and communities of exumbrella and oral arms surface could be the consequence of different surface/epithelial structures and their function. Exumbrella and oral arms are densely covered with cilia. The epidermis contains numerous mucus cells, especially in densely ciliated area [39]. Mucus cells were thought to contribute to the constantly renewing mucus layer involved in surface cleaning [40], and potentially controlling the density of associated bacteria. The exumbrella and oral arms surfaces are in constant contact with bacteria in surrounding ambient seawater, attracted by secreted mucus, which is potentially a high quality energy source and settling niche. The bacteria of genus *Phaeobacter* and genus *Ruegeria,* which belongs to *Silicibacter-Ruegeria* subgroup, are members of *Roseobacter* clade, known as the successful surface colonizers, and as the fastest utilizers of nutrients in the marine environment [80]. They produce acylated homoserine lactons (AHLs), the quorum-sensing signals involved in biofilm formation and function [80]. Besides the production of broad range biologically active metabolites, bacteria of the *Pseudoalteromanas* genus produce extracellular enzymes and exopolysaccharides, which all together enable them to successfully compete for nutrients and colonization of surfaces [79]. Bacteria of *Alteromonas* and *Vibrio* genus are widespread in the marine environment and are common surface and particle colonizers [81]. *Alteromonas* produces and secretes a variety of extracellular enzymes that contribute to the hydrolysis of biopolymers, including polysaccharides, proteins, nucleic acids, and lipids, which are the major components of marine organic particles [81]. According to Allers *et al*. [98], their versatile metabolism helps them exploit a complex substrate source, such as coral mucus, which in composition resembles to mucus produced by jellyfish *A. aurita* [43]. *Vibrio* species are major chitin utilizers, largely contributing to global carbon and nitrogen cycling. Although association with insoluble chitinous surface of detritus and life zooplankton is a preferable lifestyle for vibrios [81], they were found in association with other marine animals, including jellyfish, and there are indices that could be highly enriched in the seawater at the end of the jellyfish blooms [33]. Ritchie [90] characterized the *Vibrio* species more as ‘visitors’ than true residents, and as commensal microorganisms that can potentially become opportunistic under certain conditions. *Vibrio coralliilyticus* was found in high abundances in coral tissue slurry [9] and proven to infect and cause tissue damage in corals at higher temperatures [99]. The *Stenotrphomonas* genus was usually represented in low abundances in communities associated with marine animals [3,5,9], but found to be producing antimicrobial compounds [100]. Otherwise, *Stenotrophomonas* species are found in many environments, but mostly associated with terrestrial plants that provide plant protection and growth promotion. They were also found to be resistant to heavy metals and antibiotics, and to degrade pollutants like polycyclic aromatic hydrocarbons (PAHs) and xenobiotics [85]. In contrast, the gastral cavity is somehow isolated from the surrounding environment. The gastral cavity surface is covered by finger-like villi, with numerous cilia at their apical region, and with vesiculous receptacles at the basal region. Mucus cells in the gastrodermis are present mostly at the apical region of the villi, while gastrodermis at the basal region is composed mainly by serous cells, producing digestive enzymes [39]. As a niche, the jellyfish ‘gut’ could somehow impose strict requirements of bacteria to survive. Studies on *Cotylorhiza tuberculata* showed possible intracellular symbiotic bacteria within organs of the gastral cavity (gastric filaments), with possible involvement in the digestion process [26]. The dominance of *Betaproteobacteria* in medusa gastral cavity detected within our study is somehow surprising, since they are more characteristic for organic aggregates in limnetic ecosystems [89]. However, bacteria of the *Burkholderia* and *Achromobacter* genus were also isolated from the marine environment, including animals [83,87]. Both were found to be able to degrade PAHs and to be resistant to multiple antibiotics [82–84,86,87]. Similarly, *Achromobacter* species were found to be n-alkane degrader and to remove also anthracene, phenanthrene, and pyrence from the environment [82]. In addition, *Achromobacter* sp. HZ01 possesses genes related to the metabolism of secondary metabolites [83]. The *Cupriavidus* species were not detected in the marine environment, to our knowledge, however they were attributed with the ability to degrade aliphatic hydrocarbons [101]. Similarly, were the marine *Pseudomonas* species found to be able to degrade hydrocarbons like naphthalene, present within petroleum [86,102]. However, they were also found in association with sponges, producing antimicrobial compounds [2,100]. *Kocuria* isolated from marine sponges were found to produce the antibiotic kocurin, active against *Staphylococcus aureus* (MRSA) [103,104]. Even more so, *Kocuria* isolated from the marine environment was found to be able to utilize polyethylene as a sole carbon source [105]. The Gulf of Trieste, an ecosystem where *Aurelia* used in our study was collected, is known to be impacted by different anthropogenic pressures. Consequently, the sediment and the water column is polluted with PAHs and other chemical compounds [106] as well as by faecal bacteria, originating from coastal run off and municipal wastewater discharges [106]. This suggests that the bacterial community associated with jellyfish from this environment could be adapted to such environmental conditions. Furthermore, supporting our hypothesis, polyps generating *Aurelia* medusa were previously found attached to artificial structures such as port pillars [107]. This indicates that pollution adapted bacterial community could evolved and prosper at polyp and medusa stages. This altogether suggests possible impact of anthropogenic pollution on the structure of bacterial community associated with jellyfish and possible adaptation mechanism of jellyfish associated bacterial population.

Hosts recruit bacteria, which are beneficial for their development or contribute to their well-being. Selection of certain bacteria in different medusa body parts could be the strategy of *Aurelia*, to harbor bacteria with specific functions needed in different body parts. Bacterial associates found on *Aurelia* exumbrella and oral arms surface were previously supposed to assist in host defense against pathogens and fouling organisms from surrounding seawater [13,93,108,109]. This is not surprising, since *Vibrio* and pigmented strains of *Phaeobacter, Ruegeria*, and *Pseudoalteromonas* produce antimicrobial compounds when attached to live or inert surfaces [44,109–112]. In addition, some of extracellular compounds produced by *Pseudoalteromonas* bacteria were found to enhance the chances of host organisms to survive in specific marine habitats [79]. Apprill *et al*. [78] even proposed possible role of *Pseudoalteromonas* in coral planula settlement and adhesion process, while were the *Roseobacter* clade bacteria supposed to be important in coral development. The role in host defense was also proposed for bacteria of *Stenotrophomonas,* detected within all body part communities and for *Pseudomonas* [2,100], and *Actinobacteria* [103,113], present within the gastral cavity. However, bacteria in ‘digestive system’ should be more involved in food digestion and nutrition, which is reasonable, since jellyfish are supposed to ‘lack’ some digestive enzymes to utilize prey [114]. An especially intriguing ability of the gastral cavity bacteria is degradation of PAH’s, xenobiotics, and plastic. Jellyfish mucus was found to have structural properties to effectively accumulate nanoparticles [40] and PAHs [115]. This property could be also applied to entrap micro-plastic particles, which was demonstrated for corals [116]. PAHs were found to be highly toxic for zooplankton organisms, however, adult medusa *A. aurita* and *M. leidyi* showed a higher tolerance to exposure [117]. *A. aurita* under stress conditions, release blobs of mucus [41], detected also under exposure to crude oil (containing PAHs) [115]. This suggests that sloughing as a possible way to reduce the toxic effect of crude oil. In addition, when PAHs were entrapped within jellyfish mucus, hydrocarbon-degrading bacteria cell densities doubled, which resulted in a significant increase in oil degradation [115]. Some of toxic particles, entrapped within mucus and covered with degrading bacteria, could be also transferred by ciliary currents and boundary layer flow to a marginal umbrella groove, and then to gastral cavity, since this is one type of prey capture recognized for *A. aurita* [114], which could explain high abundances of hydrocarbon and plastic degrading bacteria found in the gut of *Aurelia* jellyfish within our study. Even more so, since toxic compounds are utilized by associated bacteria, less are accumulated within jellyfish tissue and are not transferred to higher trophic levels.

### Bacterial community structure shifts due to jellyfish *Aurelia* population collapse

The shift in bacterial community composition within a one-month period (from May to June) was observed. It resulted in a higher abundance of *Gammaproteobacteria*, especially *Vibrio*, which became a dominant member of community. This shift towards *Vibrio* was even more evident in cultural bacterial community.

The major difference between both studied months was a rise in the temperature and the viability state of *Aurelia* jellyfish in the Gulf of Trieste. In June was the end of blooming period and jellyfish were in the phase of dying, which was evident as typical signs of moribund jellyfish: degenerated tentacles, oral structures, and gonads, reduced swimming ability and necrosis of the epithelial bell tissue [118]. The process is normally triggered by environmental stress like change in temperature or salinity, food availability, parasitism, and spawning or even more likely, interacting stressors [118]. Nevertheless, in summer, the greater part of the *Aurelia* jellyfish population was found to be parasitized, along with altered morphology, growth, and swimming pattern in the Big Lake (Mljet Island, Croatia) [119]. This could indicate that jellyfish defense mechanism was probably disturbed due to environmental stress (higher temperature), which resulted in parasitism and mortality.

The senescing process in jellyfish indirectly affects interaction between jellyfish (host) and bacterial associates, which leads to a shift in the associated microbial community. Moribund jellyfish, without their own defense mechanisms, represent organic rich particles, where the structure of associated bacterial community is influenced solely by bacterium-bacterium antagonism and environmental conditions, determining the presence of metabolically active bacteria. Jellyfish tissue was found to be high quality labile organic substrate for bacteria [33-35]. Previous bacterial degradation experiments performed on *Aurelia* jellyfish in the Gulf of Trieste, resulted in the increase in *Vibrio* abundance [33].

*Vibrio* was recognized before as ‘visitor,’ exploiting the nutrient-rich niche [90]. As such, this commensal microorganism probably dwells in the ‘cloud’ of jellyfish mucus and under the right conditions becomes opportunistic. Vibrios rapidly grow in organic-rich environments [120], and together with tolerance to higher temperature [121] (documented in June in the Gulf of Trieste), up-regulating virulence determinants such as motility, resistance to antimicrobial compounds, hemolysis, and cytotoxicity detected in coral pathogens [81] and the references within), it can outcompete other bacterial residents and become highly dominant.

## Conclusion

Both culture-dependent and independent methods have been extensively used to study and to understand the role of microbial communities associated with marine animals, especially crustacean zooplankton, and benthic sponges and corals. Data available on *A. aurita* are still limited. With the exception of *Mycoplasma* bacteria, a possible endosymbiont detected within *A. aurita* tissue [18,27], the nature of the relationship between *Aurelia* jellyfish and bacterial associates is not straight forward. In addition, it is hard to say whether or not these bacteria are true residents of jellyfish, forming a species-specific association with the host or are just opportunistic microbes residing in niche of an organically-rich environment. So far, we only speculate on the role of bacterial associates, although Weiland-Bräuer *et al*. [27] suggested associated bacteria may play important functional roles during the life cycle of *A. aurita*. Bacteria associated with *Aurelia* jellyfish in the Gulf of Trieste were found to be mostly generalists, composing for the host beneficial assemblage possibly involved in food digestion and protection from toxic compounds, pathogens, and other fouling organisms. With speculation on the active and passive role of *Aurelia* jellyfish in selection of bacterial associates, demonstrated for other animals [45], jellyfish may form a relationship with diverse metabolically active microorganisms, providing more effective adaptation of host to changing environmental and anthropogenic conditions. From this perspective, the relationship somehow resembles suggested coral probiotic theory [122].

Further investigation of such a relationship is necessary to understand the relevance of the associated bacteria for the host during its life spam and during/after the bloom period, especially in areas experiencing seasonal blooms, influencing food webs, and biogeochemical cycles in those regions. Even more so, despite the small number of experimental data, our results suggest that the jellyfish - bacteria link could be applied as an effective pollution-control method in marine environments affected by crude oil and micro plastic. Finally, we would also like to emphasize the importance of culturing organisms. Although the method is biased towards certain bacterial groups, it remains important to obtain complete genome sequences, to identify properties of organisms, and to help understand the biology and ecology of microbial species.

## Acknowledgments

We would like to thank R/V Sagita crew and Dr. Mateja Grego for her help with Primer v6 analyses. We are grateful to anonymous reviewers for their critical and valuable comments on the manuscript.

## Supporting Information

**S1 Table. Composition of 16S rRNA gene clone libraries (% of clones) from samples of jellyfish exumbrella (AK), oral arms (AR) and mucus from gastral cavity (AG) and seawater samples (W) at 5m depth collected on May and June 2011 in the Gulf of Trieste.** Classification of bacterial clones was done down to the family level. The contribution of distinct bacterial families is expressed as a percentage of the total number of sequences in each sample. N is the total number of bacterial clones in the library; Sobs is the number of distinct bacterial families. In brackets beside bacterial family are the only representatives detected within the family. Numbers (N) in light grey and with asterisk (*) are total number of sequences recovered from clone library, including sequences affiliated with Chloroplast (%; presented at the bottom of table) that were omitted from further analysis.

**S2 Table. Bacterial isolates obtained from samples of jellyfish exumbrella (AK) and mucus from gastral cavity (AG) and seawater samples (W) at 5 m depth collected on May and June 2011 in the Gulf of Trieste.** Classification of bacterial isolates was done down to the genus level. The contribution of distinct bacterial genus or families is expressed as a percentage of the total number of sequences in each sample. N is the total number of isolated bacteria; Sobs is the number of distinct bacterial genus.

**S3 Table. Similarities percentage (SIMPER) analysis of 16S rRNA gene clone libraries from samples of jellyfish exumbrella (AK), oral arms (AR) and mucus from gastral cavity (AG) and seawater samples (W) collected on May and June 2011 in the Gulf of Trieste.** Seawater group (W) includes water samples collected at 5m depth on May and June.

**S4 Table. Similarities percentage (SIMPER) analysis of culturable fraction of bacterial community associated with jellyfish exumbrella (AK), mucus from gastral cavity (AG) and seawater (W) collected in May and June 2011 in the Gulf of Trieste.** Seawater group (W) includes the results of the water samples collected at 5m depth on May and June.

**S5 Table. The diversity indices S, H’, d, J’, Chao- 1 and library coverage’s (C) describing composition of total bacterial community associated with jellyfish exumbrella (AK), oral arms (AR) and mucus from gastral cavity (AG) and seawater (W) collected in May and June 2011 in the Gulf of Trieste.** S represents the number of distinct bacterial families detected in each bacterial 16S rRNA gene clone library. C is a coverage value (C = (1–n1/N), where n1 is number of phylotypes appearing only once in the library and N is the library size.

**S6 Table. The diversity indices S, H’, d, J’, Chao- 1 describing composition of culturable fraction of bacterial community associated with jellyfish exumbrella (AK) and mucus from gastral cavity (AG) and seawater (W) collected in May and June 2011 in the Gulf of Trieste.** S represents the number of distinct bacterial genus detected in each sample.

**S7 Table. Bacterial 16S rRNA sequences obtained from DGGE bands from jellyfish samples with their accession numbers.** In the table is the name and an accession number of their closest relative in GeneBank (NCBI) with % of similarity, family, taxon and isolation source.

**S8 Table. Similarities percentage (SIMPER) analysis of 16S rRNA gene clone libraries from jellyfish samples collected at the time of population peak and at the end of the bloom in the Gulf of Trieste.** Group May includes samples of jellyfish exumbrella surface (AK1, AK2) and oral arms (AR1) sampled in May. Group June includes samples of jellyfish exumbrella surface (AK6, AK7) and oral arms (AR6) collected in June.

**S9 Table. Similarities percentage (SIMPER) analysis of culturable fraction of bacterial community associated with jellyfish at the time of population peak and at the end of the bloom in the Gulf of Trieste.** Group May includes samples of jellyfish exumbrella surface (AK1, AK3, AK6) and gastral cavity (AG1, AG6) collected in May. Group June includes samples of jellyfish exumbrella surface (AK8, AK10, AK11) and gastral cavity (AG8, AG11) collected in June.

**S1 Fig. DGGE profile of bacterial 16S rRNA gene fragments of samples from *Aurelia* jellyfish exumbrella surface, oral arms and mucus from gastral cavity.** AK1, AK2: exumbrella surface of jellyfish collected in May; AK6, AK7: exumbrella surface of jellyfish collected in June; AR1: sample of oral arms of jellyfish collected in May; AR6: oral arms of jellyfish collected in June; AG1: gastral cavity mucus sample; S: standard. Numbers on the figure represent bands that were cut from the gel and successfully sequenced; color dots place sequence in one of bacterial groups.

## References

1. Apprill A. Marine Animal Microbiomes: Toward understanding host–microbiome interactions in a changing ocean. Front Mar Sci. 2017;4(July):1–9. Available from: http://journal.frontiersin.org/article/10.3389/fmars.2017.00222/full

2. Thakur NL, Anil AC. Antibacterial activity of the sponge *Ircinia ramosa*: Importance of its surface-associated bacteria. J Chem Ecol. 2000;26(1):57–71.

3. Radwan M, Hanora A, Zan J, Mohamed NM, Abo-Elmatty DM, SH Abou-El-Ela, et al. Bacterial community analyses of two Red Sea sponges. Mar Biotechnol. 2010;12:350–60.

4. Webster NS, Negri AP, Munro MMHG, Battershill CN. Diverse microbial communities inhabit Antarctic sponges. Environ Microbiol. 2004;6(3):288–300.

5. Li ZY, He LM, Wu J, Jiang Q. Bacterial community diversity associated with four marine sponges from the South China Sea based on 16S rDNA-DGGE fingerprinting. J Exp Mar Bio Ecol. 2006;329(1):75–85.

6. Hentschel U, Hopke J, Horn M, Friedrich AB, Wagner M, Hacker J, et al. Molecular Evidence for a uniform microbial community in sponges from different oceans. Appl Environ Microbiol. 2002;68(9):4431–40.

7. Kittelmann S, Harder T. Species-and site-specific bacterial communities associated with four encrusting bryozoans from the North Sea, Germany. J Exp Mar Bio Ecol. 2005;327(2):201–9.

8. Koren O, Rosenberg E. Bacteria Associated with Mucus and Tissues of the Coral *Oculina patagonica* in Summer and Winter. Appl Environ Microbiol. 2006;72(8):5254–9. Available from: http://aem.asm.org/cgi/doi/10.1128/AEM.00554-06

9. Bourne DG, Munn CB. Diversity of bacteria associated with the coral *Pocillopora damicornis* from the Great Barrier Reef. Environ Microbiol. 2005;7(8):1162–74.

10. Rohwer F, Seguritan V, Azam F, Knowlton N. Diversity and distribution of coral-associated bacteria. Mar Ecol Prog Ser. 2002;243:1–10.

11. Rohwer F, Breitbart M, Jara J, Azam F, Knowlton N. Diversity of bacteria associated with the Caribbean coral *Montastraea franksi*. Coral Reefs. 2001;20(1):85–91.

12. Webster NS, Bourne D. Bacterial community structure associated with the Antarctic soft coral, *Alcyonium antarcticum*. FEMS Microbiol Ecol. 2007;59:81–94.

13. Harder T, Lau SCK, Dobretsov S, Fang TK, Qian P-Y. A distinctive epibiotic bacterial community on the soft coral *Dendronephthya* sp. and antibacterial activity of coral tissue extracts suggest a chemical mechanism against bacterial epibiosis. FEMS Microbiol Ecol. 2003;43:337–47.

14. Tang KW, Turk V, Grossart H. Linkage between crustacean zooplankton and aquatic bacteria. Aquat Microb Ecol. 2010;61(3):261–77.

15. Gerdts G, Brandt P, Kreisel K, Boersma M, Schoo KL, Wichels A. The microbiome of North Sea copepods. Helgol Mar Res. 2013;67(4):757–73.

16. Grossart HP, Riemann L, Tang KW. Molecular and functional ecology of aquatic microbial symbionts. Front Microbiol. 2013;4(MAR):2012–3.

17. Flood PR. Architecture of, and water circulation and flow rate in, the house of the planktonic tunicate *Oikopleura labradorensis*. Mar Biol. 1991;111:95–111.

18. Daley MC, Urban-Rich J, Moisander PH. Bacterial associations with the hydromedusa Nemopsis bachei and scyphomedusa *Aurelia aurita* from the North Atlantic Ocean. Mar Biol Res. 2016;12(10):1088–100.

19. Daniels C, Breitbart M. Bacterial communities associated with the ctenophores *Mnemiopsis leidyi* and *Beroe ovata*. FEMS Microbiol Ecol. 2012;82(1):90–101.

20. Dinasquet J, Granhag L, Riemann L. Stimulated bacterioplankton growth and selection for certain bacterial taxa in the vicinity of the ctenophore *Mnemiopsis leidyi*. Front Microbiol. 2012;3(AUG):1–8.

21. Hao W. Bacterial communities associated with jellyfish. Bremen University; 2014.

22. Cleary DFR, Becking LE, Polónia ARM, Freitas RM, Gomes NCM. Jellyfish-associated bacterial communities and bacterioplankton in Indonesian Marine lakes. FEMS Microbiol Ecol. 2016;92(5):1–14.

23. Cortés-Lara S, Urdiain M, Mora-Ruiz M, Prieto L, Rosselló-Móra R. Prokaryotic microbiota in the digestive cavity of the jellyfish *Cotylorhiza tuberculata*. Syst Appl Microbiol. 2015;38(7):494–500. Available from: http://dx.doi.org/10.1016/j.syapm.2015.07.001

24. Delannoy CMJ, Houghton JDR, Fleming NEC, Ferguson HW. Mauve Stingers (*Pelagia noctiluca*) as carriers of the bacterial fi sh pathogen Tenacibaculum maritimum. Aquaculture. 2011;311(1–4):255–7. Available from: http://dx.doi.org/10.1016/j.aquaculture.2010.11.033

25. Ferguson HW, Delannoy CMJ, Hay S, Nicolson J, Sutherland D, Crumlish M. Jellyfish as vectors of bacterial disease for farmed salmon (*Salmo salar*). J Vet Diagn Invest. 2010;22:376–82.

26. Viver T, Orellana LH, Hatt JK, Urdiain M, Díaz S, Richter M, et al. The low diverse gastric microbiome of the jellyfish *Cotylorhiza tuberculata* is dominated by four novel taxa. Environ Microbiol. 2017;19(8):3039–58.

27. Weiland-Bräuer N, Neulinger SC, Pinnow N, Künzel S, Baines JF, Schmitz RA. Composition of bacterial communities associated with *Aurelia aurita* changes with compartment, life stage, and population. Appl Environ Microbiol. 2015;81(17).

28. Schuett C, Doepke H. Endobiotic bacteria and their pathogenic potential in cnidarian tentacles. Helgol Mar Res. 2010;64:205–12.

29. Riemann L, Titelman J, Båmstedt U. Links between jellyfish and microbes in a jellyfish dominated fjord. Mar Ecol Prog Ser. 2006;325(2003):29–42.

30. Manzari C, Fosso B, Marzano M, Annese A, Caprioli R, D’Erchia AM, et al. The influence of invasive jellyfish blooms on the aquatic microbiome in a coastal lagoon (Varano, SE Italy) detected by an Illumina-based deep sequencing strategy. Biol Invasions. 2015;17(3):923–40.

31. Condon RH, Steinberg DK, del Giorgio PA, Bouvier TC, Bronk DA, Graham WM, et al. Jellyfish blooms result in a major microbial respiratory sink of carbon in marine systems. Proc Natl Acad Sci. 2011;108(25):10225–30. Available from: http://www.pnas.org/cgi/doi/10.1073/pnas.1015782108

32. Blanchet M, Pringault O, Bouvy M, Catala P, Oriol L, Caparros J, et al. Changes in bacterial community metabolism and composition during the degradation of dissolved organic matter from the jellyfish *Aurelia aurita* in a Mediterranean coastal lagoon. Environ Sci Pollut Res. 2015; 22, 18: 13638–13653.

33. Tinta T, Kogovšek T, Malej A, Turk V. Jellyfish modulate bacterial dynamic and community structure. PLoS One. 2012;7(6):1–11.

34. Tinta T, Kogovšek T, Turk V, Shiganova TA, Mikaelyan AS, Malej A. Microbial transformation of jellyfish organic matter affects the nitrogen cycle in the marine water column -A Black Sea case study. J Exp Mar Bio Ecol. 2016;475:19–30.

35. Tinta T, Malej A, Kos M, Turk V. Degradation of the Adriatic medusa *Aurelia* sp. by ambient bacteria. Hydrobiologia. 2010;645:179–91.

36. Titelman J, Riemann L, Sørnes TA, Nilsen T, Griekspoor P, Båmstedt U. Turnover of dead jellyfish?: stimulation and retardation of microbial activity. Mar Ecol Prog Ser. 2006;325:43–58.

37. Pitt KA, Welsh DT, Condon RH. Influence of jellyfish blooms on carbon, nitrogen and phosphorus cycling and plankton production. Hydrobiologia. 2009;616:133–49.

38. Bosch TCG. Cnidarian-microbe interactions and the origin of innate immunity in Metazoans. Annu Rev Microbiol. 2013;67(1):499–518. Available from: http://www.annualreviews.org/doi/10.1146/annurev-micro-092412-155626

39. Heeger T, Möller H. Ultrastructural observations on prey capture and digestion in the scyphomedusa *Aurelia aurita*. Mar Biol. 1987;96(3):391–400.

40. Patwa A, Thiéry A, Lombard F, Lilley MKS, Boisset C, Bramard J, et al. Accumulation of nanoparticles in “jellyfish” mucus: a bio-inspired route to decontamination of nano-waste. Nat Sci Reports. 2015;1–8. Available from: http://dx.doi.org/10.1038/srep11387

41. Shanks A, Graham W. Chemical defense in a scyphomedusa. Mar Ecol Prog Ser. 1988;45:81–6.

42. Ovchinnikova T V., Balandin S V., Aleshina GM, Tagaev A a., Leonova YF, Krasnodembsky ED, et al. Aurelin, a novel antimicrobial peptide from jellyfish *Aurelia aurita* with structural features of defensins and channel-blocking toxins. Biochem Biophys Res Commun. 2006;348(2):514–23.

43. Ducklow HW, Mitchell R. Composition of mucus released by coral reef coelenterates. Limnol Oceanogr. 1979;24(4):706–14.

44. Long RA, Azam F. Antagonistic interactions among marine pelagic bacteria. Appl Environ Microbiol. 2001;67(11):4975–83.

45. Wahl M, Goecke F, Labes A, Dobretsov S, Weinberger F. The second skin: ecological role of epibiotic biofilms on marine organisms. Front Microbiol. 2012;3:1–21.

46. Ramšak A, Stopar K, Malej A. Comparative phylogeography of meroplanktonic species, *Aurelia* spp. and Rhizostoma pulmo (Cnidaria: Scyphozoa) in European Seas. Hydrobiologia. 2012;690(1):69–80.

47. Scorrano S, Aglieri G, Boero F, Dawson MN, Piraino S. Unmasking *Aurelia* species in the Mediterranean Sea: An integrative morphometric and molecular approach. Zool J Linn Soc. 2017;180(2):243–67.

48. Kogovšek T, Bogunović B, Malej A. Recurrence of bloom-forming scyphomedusae: Wavelet analysis of a 200-year time series. Hydrobiologia. 2010;645(1):81–96.

49. Malej A, Kogovšek T, Ramšak A, Catenacci L. Blooms and population dynamics of moon jellyfish in the northern Adriatic. Cah Biol Mar. 2012;53(3):337–42.

50. Kogovšek T, Klun K, Ikeda H, Tinta T, Uye S. Starvation -an important factor controlling scyphozoan population?. In: Fifth International Jellyfish Bloom Symposium?: Abstract book. Barcelona: Barcelona University. 2016; 2016. p. 33.

51. Brotz L, Cheung WWL, Kleisner K, Pakhomov E, Pauly D. Increasing jellyfish populations: Trends in Large Marine Ecosystems. Hydrobiologia. 2012;690(1):3–20.

52. Condon RH, Graham WM, Duarte CM, Pitt KA, Lucas CH, Haddock SHD, et al. Questioning the rise of gelatinous zooplankton in the World’s Oceans. Bioscience. 2012;62(2):160–9. Available from: http://bioscience.oxfordjournals.org/content/62/2/160.abstract

53. Condon RH, Duarte CM, Pitt KA, Robinson KL, Lucas CH, Sutherland KR, et al. Recurrent jellyfish blooms are a consequence of global oscillations. PNAS. 2013;110(3):1000–5.

54. Purcell JE, Uye S, Lo W. Anthropogenic causes of jellyfish blooms and their direct consequences for humans: a review. Mar Ecol Prog Ser. 2007;350:153–74.

55. Graham WM, Gelcich S, Robinson KL, Duarte CM, Brotz L, Purcell JE, et al. Linking human well-being and jellyfish: Ecosystem services, impacts, and societal responses. Front Ecol Environ. 2014;12(9):515–23.

56. Richardson AJ, Bakun A, Hays GC, Gibbons MJ. The jellyfish joyride: causes, consequences and management responses to a more gelatinous future. Trends Ecol Evol. 2009;24(6):312–22. Available from: http://linkinghub.elsevier.com/retrieve/pii/S0169534709000883

57. Purcell JE. Jellyfish and ctenophore blooms coincide with human proliferations and environmental perturbations. Ann Rev Mar Sci. 2012;4(1):209–35. Available from: http://www.annualreviews.org/doi/abs/10.1146/annurev-marine-120709-142751

58. Vodopivec M, Peliz AJ, Malej A. Offshore marine constructions as propagators of moon jellyfish dispersal. Environ Res Lett. 2017;12(8).

59. Kogovšek T, Vodopivec M, Raicich F, Uye S, Malej A. Comparative analysis of the ecosystems in the northern Adriatic Sea and the Inland Sea of Japan: Can anthropogenic pressures disclose jellyfish outbreaks? Sci Total Environ [Internet]. 2018;626:982–94. Available from: http://www.sciencedirect.com/science/article/pii/S0048969718300111

60. Malačič V, Petelin B. Gulf of Trieste. In: Cushman-Roisin B, Gacic M, Poulain P-M, Artegiani A, editors. Physical oceanography of the Adriatic Sea: past, present and future. Dordrecht: Kluwer Academic Press; 2001. p. 167–77.

61. Zobell CE. Marine microbiology. Marine microbiology. A monograph on hydrobacteriology. Chronica Botanica.; 1946. 240 p.

62. Giraffa G, Rossetti L, Neviani E. An evaluation of chelex-based DNA purification protocols for the typing of lactic acid bacteria. J Microbiol Methods. 2000;42(2):175–84.

63. Boström KH, Simu K, Hagström Å, Riemann L. Optimization of DNA extraction for quantitative marine bacterioplankton community analysis. Limnol Oceanogr Methods. 2004;2(1988):365–73.

64. Muyzer G, De Waal EC, Uitterlinden AG. Profiling of complex microbial populations by denaturing gradient gel electrophoresis analysis of polymerase chain reaction-amplified genes coding for 16S rRNA. Appl Environ Microbiol. 1993;59(3):695–700.

65. Muyzer G, Smalla K. Application of denaturing gradient gel electrophoresis (DGGE) and temperature gradient gel electrophoresis (TGGE) in microbial ecology. Antonie Van Leeuwenhoek. 1998;73:127–41.

66. Don RH, Cox PT, Wainwright BJ, Baker K, Mattick JS. “Touchdown” PCR to circumvent spurious priming during gene amplification. Nucleic Acids Res. 1991;19(14):4008.

67. Dar SA, Kuenen JG, Muyzer G. Nested PCR-Denaturing Gradient Gel Electrophoresis approach to determine the diversity of sulfate-reducing bacteria in complex microbial communities. Appl Environ Microbiol. 2005;71(5):2325–30.

68. Giloteaux L, Gõni-Urriza M, Duran R. Nested PCR and new primers for analysis of sulfate-reducing bacteria in low-cell-biomass environments. Appl Environ Microbiol. 2010;76(9):2856–65.

69. Schloss PD, Westcott SL, Ryabin T, Hall JR, Hartmann M, Hollister EB, et al. Introducing mothur: Open-source, platform-independent, community-supported software for describing and comparing microbial communities. Appl Environ Microbiol. 2009;75(23):7537–41.

70. Good IJ. No Title. Biometrika. 1953;40(3/4):237–64.

71. Clarke KR, Gorley RN. PRIMER v6: Manual/Tutorial. Prim Plymouth. 2006;

72. Hammer Ø, Harper DAT, Ryan PD. PAST-PAlaeontological STatistics, ver. 1.89. Palaeontol Electron. 2001;4(1):1–9.

73. Taylor MW, Schupp PJ, Dahllöf I, Kjelleberg S, Steinberg PD. Host specifity in marine sponge-associated bacteria. Environ Microbiol. 2004;6(2):121–30.

74. Cooney RP, Pantos O, Tissier MDA Le, Barer MR, Donnell AGO, Bythell JC. Characterization of the bacterial consortium associated with black band disease in coral using molecular microbiological techniques. Environ Microbiol. 2002;4(7):401–13.

75. Bythell JC, Barer MR, Cooney RP, Guest JR, O’Donnell AG, Pantos O, et al. Histopathological methods for the investigation of microbial communities associated with disease lesions in reef corals. Lett Appl Microbiol. 2002;34:359–64.

76. Johnston IS, Rohwer F. Microbial landscapes on the outer tissue surfaces of the reef-building coral *Porites compressa*. Coral Reefs. 2007;26(2):375–83.

77. Garren M, Azam F. Corals shed bacteria as a potential mechanism of resilience to organic matter enrichment. ISME J [Internet]. 2012;6:1159–65. Available from: http://dx.doi.org/10.1038/ismej.2011.180

78. Apprill A, Marlow HQ, Martindale MQ, Rappé MS. Specificity of associations between bacteria and the coral *Pocillopora meandrina* during early development. Appl Environ Microbiol. 2012;78(20):7467–75.

79. Holmström C, Kjelleberg S. Marine P*seudoalteromonas* species are associated with higher organisms and produce biologically active extracellular agents. FEMS Microbiol Ecol. 1999;30:285–93.

80. Dang H, Li T, Chen M, Huang G. Cross-Ocean Distribution of *Rhodobacterales* bacteria as primary surface colonizers in temperate coastal marine waters. Appl Environ Microbiol. 2008;74(1):52–60.

81. Dang H, Lovell CR. Microbial surface colonization and biofilm development in marine environments. Microbiol Mol Biol Rev. 2016;80(1):91–138.

82. Deng M-C, Li J, Liang F-R, Yi M, Xu X-M, Yuan J-P, et al. Isolation and characterization of a novel hydrocarbon-degrading bacterium *Achromobacter* sp. HZ01 from the crude oil-contaminated seawater at the Daya Bay, southern China. Mar Pollut Bull. 2014;83(1):79–86. http://www.sciencedirect.com/science/article/pii/S0025326X14002306

83. Hong YH, Ye CC, Zhou QZ, Wu XY, Yuan JP, Peng J, et al. Genome sequencing reveals the potential of *Achromobacter* sp. HZ01 for bioremediation. Front Microbiol. 2017;8:1–14.

84. Juhasz AL, Britz ML, Stanley GA. Degradation of fluoranthene, pyrene, benz [a] anthracene and dibenz [a,h] anthracene by *Burkholderia cepacia*. J Appl Microbiol. 1997;83(2):189–98.

85. Ryan RP, Monchy S, Cardinale M, Taghavi S, Crossman L, Avison MB, et al. The versatility and adaptation of bacteria from the genus *Stenotrophomonas*. Nat Rev Microbiol. 2009;7:514–25. http://dx.doi.org/10.1038/nrmicro2163

86. Harayama S, Kishira H, Kasai Y, Shutsubo K. Petroleum biodegradation in marine environments. J Mol Microbiol Biotechnol. 1999;1(1):63–70.

87. Maravić A, Skočibušić M, Šprung M, Šamanić I, Puizina J, Pavela-Vrančić M. Occurrence and antibiotic susceptibility profiles of *Burkholderia cepacia* complex in coastal marine environment. Int J Environ Health Res. 2012;22(6):531–42. http://www.tandfonline.com/doi/abs/10.1080/09603123.2012.667797

88. DeLong EF, Franks DG, Alldredge AL. Phylogenetic diversity of aggregate-attached marine bacterial assemblages. Limnol Oceanogr. 1993;38(5):924–34.

89. Simon M, Grossart H, Schweitzer B, Ploug H. Microbial ecology of organic aggregaes in aquatic ecosystems. Aquat Microb Ecol. 2002;28:175–211.

90. Ritchie KB. Regulation of microbial populations by coral surface mucus and mucus-associated bacteria. Mar Ecol Prog Ser. 2006;322:1–14.

91. Daniels CA, Zeifman A, Heym K, Ritchie KB, Watson CA, Berzins I, et al. Spatial heterogeneity of bacterial communities in the mucus of *Montastraea annularis*. Mar Ecol Prog Ser. 2011;426:29–40.

92. Lampert Y, Kelman D, Dubinsky Z, Nitzan Y, Hill RT. Diversity of culturable bacteria in the mucus of the Red Sea coral *Fungia scutaria*. FEMS Microbiol Ecol. 2006;58(1):99–108.

93. Shnit-Orland M, Kushmaro A. Coral mucus-associated bacteria: a possible first line of defense. FEMS Microbiol Ecol. 2009;67:371–80.

94. Hao W, Gerdts G, Peplies J, Wichels A. Bacterial communities associated with four ctenophore genera from the German Bight (North Sea). FEMS Microbiol Ecol. 2015;(October 2014):1–11. http://femsec.oxfordjournals.org/cgi/doi/10.1093/femsec/fiu006

95. Kirchman DL, Dittel AI, Malmstrom RR, Cottrell MT. Biogeography of major bacterial groups in the Delaware Estuary. Limnol Oceanogr. 2005;50(5):1697–706.

96. Webster NS, Wilson KJ, Blackall LL, Hill RT. Phylogenetic diversity of bacteria associated with the marine sponge *Rhopaloeides odorabile*. Appl Environ Microbiol. 2001;67(1):434–44.

97. Franzenburg S, Walter J, Künzel S, Wang J, Baines JF, Bosch TCG. Distinct antimicrobial peptide expression determines host species-specific bacterial associations. Proc Natl Acad Sci U S A. 2013;110(39):E3730–8.

98. Allers E, Niesner C, Wild C, Pernthaler J. Microbes enriched in seawater after addition of coral mucus. Appl Environ Microbiol. 2008;74(10):3274–8.

99. Ben-Haim Y, Thompson FL, Thompson CC, Cnockaert MC, Hoste B, Swings J, et al. *Vibrio coralliilyticus* sp. nov., a temperature-dependent pathogen of the coral *Pocillopora damicornis*. Int J Syst Evol Microbiol. 2003;53:309–15.

100. Santos OCS, Pontes PVML, Santos JFM, Muricy G, Giambiagi-deMarval M, Laport MS. Isolation, characterization and phylogeny of sponge-associated bacteria with antimicrobial activities from Brazil. Res Microbiol. 2010;161:604–12

101. Bacosa HP, Suto K, Inoue C. Bacterial community dynamics during the preferential degradation of aromatic hydrocarbons by a microbial consortium. Int Biodeterior Biodegrad. 2012;74:109–15. Available from: http://dx.doi.org/10.1016/j.ibiod.2012.04.022

102. Prince RC. Petroleum spill bioremediation in marine environments. Crit Rev Microbiol. 1993;19(4):217–40.

103. Abdelmohsen UR, Bayer K, Hentschel U. Diversity, abundance and natural products of marine sponge-associated actinomycetes. Nat Prod Rep. 2014;31(3):381–99. Available from: http://xlink.rsc.org/?DOI=C3NP70111E

104. Palomo S, González I, De La Cruz M, Martín J, Tormo JR, Anderson M, et al. Sponge-derived *Kocuria* and *Micrococcus* spp. as sources of the new thiazolyl peptide antibiotic kocurin. Mar Drugs. 2013;11(4):1071–86.

105. Harshvardhan K, Jha B. Biodegradation of low-density polyethylene by marine bacteria from pelagic water, Arabian Sea, India. Mar Pollut Bull. 2013;77:100–6.

106. Turk V, Mozetič P, Malej A. Overview of eutrophication-related events and other irregular episodes in Slovenian Sea (Gulf of Trieste, Adriatic Sea). Annales Ser hist nat 2007;17:197–216.

107. Hočevar S, Malej A, Boldin B, Purcell JE. Seasonal fluctuations in ppulaton dynamics of aurelia aurita polyp in situ with a modelling perspective. Mar Ecol Prog Ser. 2018; 591:155–166.

108. Dobretsov S, Qian P-Y. The role of epibotic bacteria from the surface of the soft coral *Dendronephthya* sp. in the inhibition of larval settlement. J Expr Mar Biol Ecol. 2004;299:35–50.

109. Holmström C, Egan S, Franks A, McCloy S, Kjelleberg S. Antifouling activities expressed by marine surface associated *Pseudoalteromonas* species. FEMS Microbiol Ecol. 2002;41:47–58.

110. Bruhn JB, Nielsen KF, Hjelm M, Hansen M, Schulz S, Gram L, et al. Ecology, inhibitory activity, and morphogenesis of a marine antagonistic bacterium belonging to the *Roseobacter* clade. Appl Environ Microbiol. 2005;71(11):7263–70.

111. Gram L, Melchiorsen J, Bruhn JB. Antibacterial activity of marine culturable bacteria collected from a global sampling of Ocean surface waters and surface swabs of marine organisms. Mar Biotechnol. 2010;12:439–51.

112. Porsby CH, Nielsen KF, Gram L. *Phaeobacter* and *Ruegeria* Species of the *Roseobacter* clade colonize separate niches in a Danish Turbot (*Scophthalmus maximus*)-rearing farm and antagonize *Vibrio anguillarum* under different growth conditions. Appl Environ Microbiol. 2008;74(23):7356–64.

113. Selvin J, Joseph S, Asha KRT, Manjusha W a., Sangeetha VS, Jayaseema DM, et al. Antibacterial potential of antagonistic *Streptomyces* sp. isolated from marine sponge *Dendrilla nigra*. FEMS Microbiol Ecol. 2004;50(2):117–22.

114. Arai MN. A Functional Biology of Scyphozoa. First edit. London: Chapman & Hall; 1997. 316 p.

115. Gemmell BJ, Bacosa HP, Liu Z, Buskey EJ. Can gelatinous zooplankton influence the fate of crude oil in marine environments? Mar Pollut Bull [Internet]. 2016;113(1–2):483–7. Available from: http://dx.doi.org/10.1016/j.marpolbul.2016.08.065

116. Hall NM, Berry KLE, Rintoul L, Hoogenboom MO. Microplastic ingestion by scleractinian corals. Mar Biol. 2015;162(3):725–32.

117. Almeda R, Wambaugh Z, Chai C, Wang Z, Liu Z, Buskey EJ. Effects of Crude Oil Exposure on bioaccumulation of polycyclic aromatic hydrocarbons and survival of adult and larval stages of gelatinous zooplankton. PLoS One. 2013;8(10):20–1.

118. Pitt KA, Chelsky Budarf A, Browne JG, Condon RH. Bloom and Bust: Why do blooms of jellyfish collapse? In: Pitt KA, Lucas CH, editors. Jellyfish Blooms. Netherlands: Springer; 2014. p. 79–103.

119. Graham WM, Chiaverano L, D’ambra I, Mianzan H, Colombo GA, Acha EM, et al. Fish and jellyfish: using the isolated marine “lakes” of Mljet Island, Croatia, to explore larger marine ecosystem complexities and ecosystem-based management approaches. In: Annales series historia naturalis. 2009. p. 39–48.

120. Eilers H, Pernthaler J, Amann R. Succession of pelagic marine bacteria during enrichment: A close look at cultivation-induced shifts. Appl Environ Microbiol. 2000;66(11):4634–40.

121. Thompson FL, Iida T, Swings J. Biodiversity of *Vibrios*. Microbiol Mol Biol Rev. 2004;68(3):403–31.

122. Reshef L, Koren O, Loya Y, Zilber-Rosenberg I, Rosenberg E. The coral probiotic hypothesis. Environ Microbiol. 2006;8(12):2068–73.

